# Efficient repair of human homozygous genetic mutation by CRISPR/Cas9 mediated interlocus gene conversion

**DOI:** 10.1101/2022.09.05.506576

**Authors:** Fei Yang, Jingtao Pang, Guolong Liu, Yang Yang, Shenguang Qin, Ying Zhang, Yongrong Lai, Dan Liang, Yuxuan Wu

## Abstract

DNA double-strand breaks (DSBs) induced by gene editing tools are primarily resolved either by non-homologous end joining (NHEJ) or homology-directed repair (HDR) using exogenous synthetic DNA templates. Repaired by error-prone NHEJ may lead to unexpected indels at the targeted site. In the case of most genetic disorders, HDR-mediated precise correction using an exogenous homologous sequence is ideal. However, the therapeutic application of HDR might be especially challenging given the requirement for the codelivery of exogenous DNA templates with toxicity into cells, and the low efficiency of HDR could also limit its clinical application. Here, we used hematopoietic stem cells (HSCs) with genetic mutations to cause β-thalassemia in *HBB* coding regions and discovered that many cells are actually repaired by CRISPR/Cas9-mediated gene conversion (GC) independent of exogenous synthetic DNA templates. We show that pathogenic mutations in the *HBB c*oding regions of HSCs can be repaired efficiently through CRISPR/GC using the paralog gene *HBD* as the internal template. Electroporations of Cas9 for ribonucleoprotein with sgRNA into haematopoietic stem and progenitor cells (HSPCs) with a variety of pathogenic gene mutations also resulted in effective conversion of mutations to normal wild-type sequences without exogenous DNA template. Moreover, the edited HSCs can repopulate the haematopoietic system and generate erythroid cells with a greatly reduced propensity for thalassemia after transplantations. Thus, CRISPR/GC, independent of exogenous DNA templates, holds great promise for gene therapy of genetic diseases.

## Introduction

A variety of genetic diseases have been cured by highly efficient gene editing^1-3^. For instance, delivery to hematopoietic stem cells (HSCs) of Cas9-sgRNA ribonucleoprotein (RNP) complexes could plausibly contribute to the healing of hematonosis^4-10^. Novel therapeutic options have been explored, such as fetal hemoglobin (HbF) induction by disrupting the enhancer of *BCL11A* (γ-globin gene repressor)^11-19^, which is a universal strategy to ameliorate β-thalassemia and sickle cell disease (SCD). Other tactics about gene editing intervention for β-thalassemia are focusing on editing of the intron mutations (e.g., *IVS2-654* or *IVS1-110*) ^20, 21^ directly due to aberrant splice sites. Although these approaches show great promise, they are still under deep experimental development, with limited clinical trials^22, 23^.

Applications of gene editing ultimately depend on DNA repair triggered by targeted DNA double-strand breaks (DSBs). DNA DSBs induced by gene editing are typically repaired through homology-directed repair (HDR) and non-homologous end joining (NHEJ). Repaired by error-prone NHEJ may lead to unexpected insertion or deletion at the targeted site. In the case of most genetic blood disorders, HDR-mediated precise correction of disease-causing mutations using an exogenous homologous sequence as a template in repopulating HSCs is ideal. However, the therapeutic application of HDR might be especially challenging given the requirement for the codelivery of exogenous donor DNA templates with toxicity into cells, and the low efficiency of HDR in ex vivo gene editing of HSCs could also limit the clinical application of this approach^*24*^.

Gene conversion (GC) events, a combination of natural selection^25-27^, can be easily overlooked. Alternatively, it has also been previously described as one of the homologous recombination mechanisms, and independent of exogenous templates, usually initiated as a DSB repair response, it can occur between homologous chromosomes or highly similar sequences. Some research has observed high rates of GC despite sequence differentiation between species, especially in inverted regions^25^. Indeed, the sequences of *HBG2* and *HBG1* led to the first description of GC in the human genome^28^. Moreover, the human β-globin cluster includes five paralogs (*HBE, HBG2, HBG1, HBD* and *HBB)* located together on chromosome 11. Human *HBB*, which encodes the major β-globin chain of adult hemoglobin (HbA), shows high homology with the neighbor gene *HBD* ^*28-30*^. GC events between them have been reported very frequently^25^,^29^.

β^0^ 4142-thalassemia is a premature termination caused by a frameshift mutation of 4 base pair (-TCTT) deletion in the β-globin gene *HBB*^31^. This mutation causes the introduction of a stop codon, resulting in a truncated nonfunctional β-globin. Here, we investigated the restoration of *HBB-4142* and some other genetic mutations causing thalassemia in *HBB* coding region*s to identify potential CRISPR/GC mechanisms and whether GC is able to mitigate mutations in the β-globin gene HBB*. In this study, we show that β^0^ 4142 in HSCs can be repaired efficiently and persistently through CRISPR/GC using the paralog gene *HBD* as the internal template. Ex vivo electroporation of Cas9 with three nuclear localization sequences (3NLS-Cas9) for ribonucleoprotein (RNP) with a targeting guide RNA into hematopoietic stem and progenitor cells (HSPCs) from homozygous donors resulted in 27% conversion (over 16% in heterozygous) of β^0^ 4142 to normal gene sequences. Sixteen weeks post-transplantation of edited human HSPCs into immunodeficient mice, we found more than 10% normal wild-type cells in recipient mouse bone marrow with multilineage-repopulating self-renewing human HSCs. This suggests durable gene editing and the induction of *HBB* to potentially therapeutic levels in erythroid progeny generated in vitro and in vivo after transplantation of HSPC mouse models. Moreover, efficient editing could be achieved with GC to correct multiple types of mutations in the *HBB* coding region. *Thus, our findings suggest that CRISPR/Cas9-mediated GC offers a potential approach used as an autologous treatment for β-thalassemia that eliminates pathogenic HBB* mutations. Especially *in vivo* gene editing, the delivery of foreign DNA templates into cells is extremely challenging, and it might also stimulate innate immunity in targeted cells, leading to cell death. CRISPR/GC, independent of exogenous DNA templates, holds great promise for gene therapy of multiple genetic diseases.

## Result

### Experimental strategy for CRISPR/Cas9 correction of the thalassemia β^0^4142 mutation

The β^0^ 4142 mutation can be based on the design of a CRISPR/Cas9 strategy to recognize an NGG protospacer-adjacent motif causing the formation of a DSB, inducing the cell to activate homologous recombination mechanisms^31, 32^. Meanwhile, we paired the Cas9 RNP with single-stranded DNA oligonucleotides (ssODNs) to correct mutations with wild-type gene sequences through a precise HDR mechanism. According to our previous work, we investigated maximizing editing efficiency in HSCs^18^, and designed seven β^0^ 4142-targeting single guide RNAs (sgRNAs) codelivered with the 3NLS Cas9 protein **(Fig. S1)** to CD34^+^ HSPCs of thsalassemia patient Donor #1, who carried a homozygous mutation at the CD4142 site on the *HBB* gene **(Fig. 1a and b; see details in Table S1 and S2)**. Based on the maximally active sgRNAs compatible with editing and position, sgRNA-1 was selected for subsequent experiments (**Fig. 1c**). Therefore, 3NLS Cas9 protein with sgRNA-1 and various concentrations of ssODNs were codelivered into CD34^+^ HSPCs of homozygous Donor #1 and compound heterozygous Donor #2 (see details in **Table S2**), respectively. We quantified the frequency of HDR and indel rates at the mutation site by next-generation sequencing (NGS) of PCR amplicons derived from genomic DNA extracted from pools of edited cells. Notably, we found highly efficient restoration in each group, especially by using 150 μM ssODN as a donor template to repair the damage **(Fig. 1d; see details in Table S1)**. Next, we compared the recombination of our strategy to previous studies at the *HBB* locus in CD34^+^ cells. The restoration efficiency at β^0^ 4142 was significantly higher than that of other reported pathogens^33, 34^ with DNA templates. Moreover, according to the percentage of HbA with respect to total hemoglobin obtained after CRISPR/Cas9 correction, we found that β-globin mRNA levels in patient CD34^+^ HSPCs were significantly elevated (mean 40.8% of β-globin, range 30.3% to 60.1%, compared to 1.7% in unedited cells) **(Fig. 1e)**. Incredibly, compared with non-edited cells, there is a greater probability of restoration without exogenous donors only through the destruction of Cas9 protein mixed with sgRNA-1, which achieved approximately 27% restoration in β^0^ 4142 homozygotes (16% in heterozygotes) from mutation to wild type **(Fig. 1d)**, indicating CRISPR/Cas9-mediated GC.

**Fig. 1.**
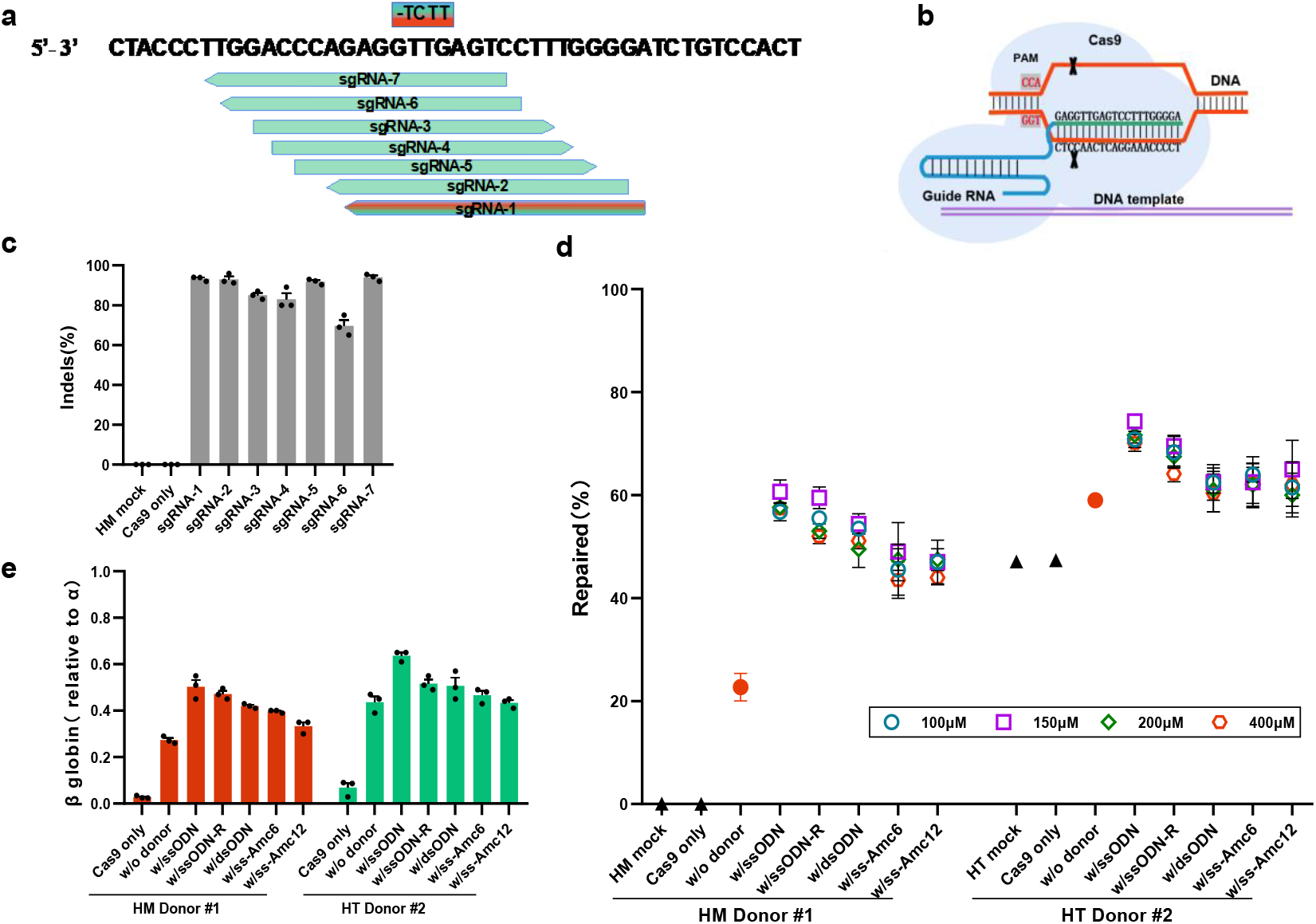
Identification of efficient β^0^ 4142 guide RNAs with donor templates for the restoration and amelioration of β-thalassemia. **a**, Seven sgRNAs targeting the β^0^ 4142 core marked with green arrows. CD4142 deletion base pairs marked with chromatic rectangles. **b**, Schematic diagram of CRISPR/Cas9 targeting sites in the mutant *HBB* gene. Blue lines label sgRNA sequences; purple lines label exogenous DNA templates. **c**, Editing efficiency of Cas9 RNPs coupled with the various sgRNAs in CD34^+^ HSPCs from homozygotes measured by TIDE analysis. HM mock represents only CD34^+^ HSPCs from homozygous Donor #1; Cas9 only represents CD34^+^ HSPCs from homozygous Donor #1 electroporated with Cas9 only. Error bars indicate the standard deviation (*n*= 3 replicates); donors edited without sgRNA were plotted as a negative control. **d**, CD34^+^ cells from patients #1 and #2 were electroporated with Cas9 coupled with sgRNA-1 and with different concentration gradient DNA templates. HM mock, HSPCs of homozygous Donor #1 only; HT mock, HSPCs of heterozygote Donor #2 only; w/o donor, patient donors edited with RNPs but without DNA templates; Forward and reverse normal gene sequences were synthesized expressed by w/ssODN and w/ssODN-R. These two single-stranded DNAs were incorporated into dsODNs annealed by PCR. Primers with 5′ modifications included an amine group with a C6 linker (AmC6) or C12 linker (AmC12) expressed by w/ss-AmC6, w/ss-AmC12. **e**, β globin expression by RT–qPCR analysis in erythroid cells in vitro differentiated from RNP-edited CD34^+^ HSPCs. Data are plotted as the mean ± s.d. and analyzed using unpaired two-tailed Student’s *t* tests. Data are representative of three biologically independent replicates.

### GC-mediated β^0^ 4142 corrections independent of exogenous DNA templates

To confirm the authenticity of restoration and CRISPR/Cas9-mediated GC, we further edited the β^0^ 4142 homozygous cells (Donor #1) under the same condition of only Cas9 protein and all the above sgRNAs. Compared with the mock and Cas9-only groups, we also found inverse restorations in the β^0^ 4142 mutant site, and the efficiency was positively correlated with the distance between the sgRNA cutting position and mutant site **(Fig. 2a)**. As expected, the β-globin mRNA levels of all edited groups also showed a certain improvement **(Fig. 2b)**.

**Fig. 2.**
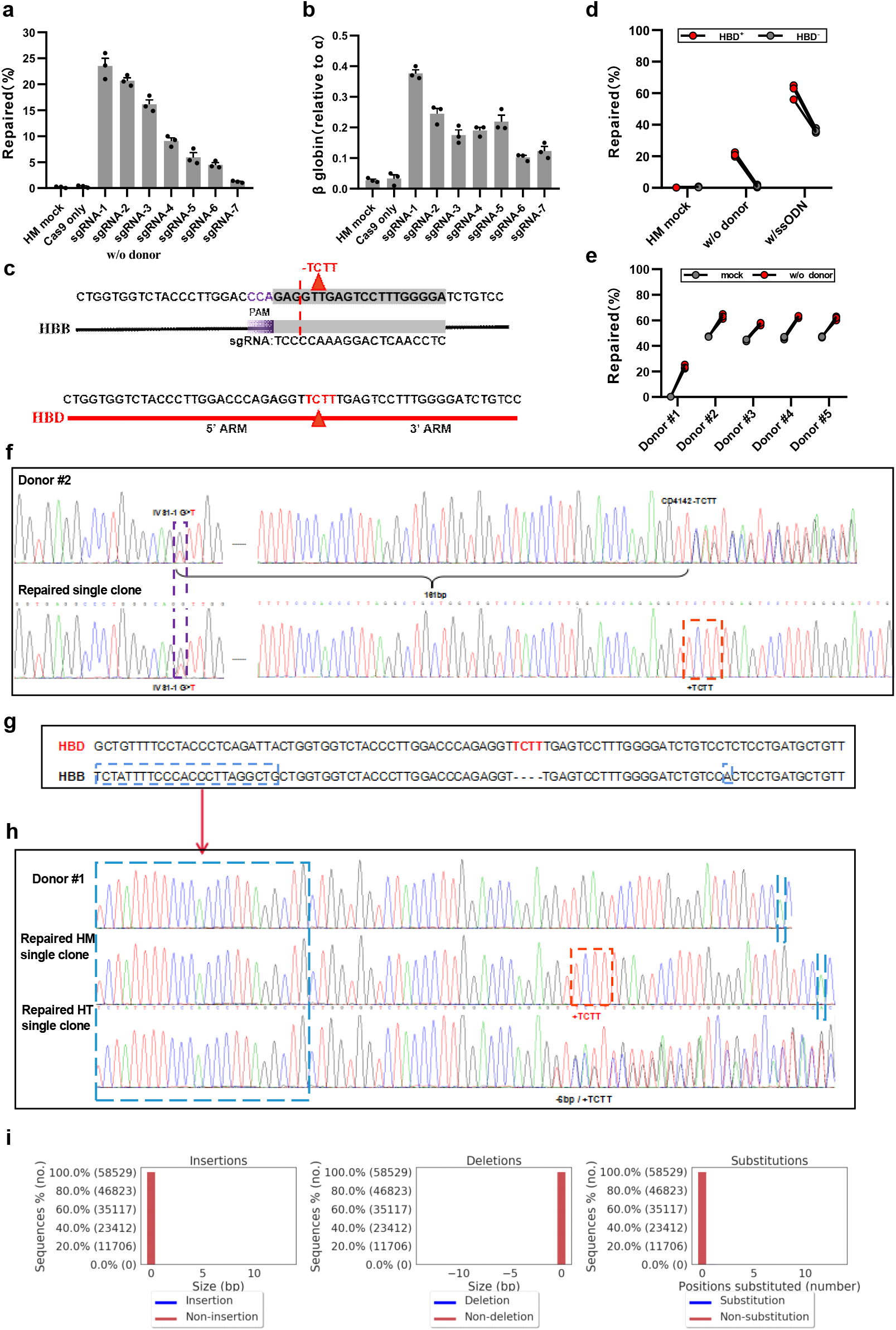
GC-mediated correction of β^0^ 4142 deletions independent of exogenous DNA temples. **a**, Efficiency of restoration edited by Cas9 coupled with the various sgRNAs in homozygous β^0^ 4142 CD34^+^ HSPCs measured by deep-seq analysis. Error bars indicate the standard deviation (*n*=3 replicates). **b**, β-globin expression by RT–qPCR analysis in erythroid cells in vitro differentiated from RNP-edited CD34^+^ HSPCs. **c**, The homologous genomic sequence of *HBB* and *HBD*, Red triangle represents the deletion base pairs, the color purple marked a NGG protospacer-adjacent motif, gray marked sgRNA-1.**d**, Restoration of the β^0^ 4142 in *HBD-4142* KO donor determined by deep-seq analysis versus edited only by the RNP complex. Gray points *HBD*^*-*^ represent samples with KO of *HBD-4142* edited by Cas9 coupled with sgRNA-1 72 hours later, and red points represent samples edited only with Cas9 coupled with sgRNA-1. **e**, Efficiency of restorations as measured by deep-seq analysis of Cas9:sgRNA-1 RNP targeting the β^0^ 4142 site in CD34^+^ HSPCs from patient donors (see details in Table S2). **f–h**, Genotyping of erythroid cells derived from single colonies screened out edited CD34^+^ HSPCs from Donors #1 and #2. The purple dashed line in **f** highlights IVS1-1 G>T, corrected deletions of β^0^ 4142 (orange dashed line), specific fragments or base pairs of *HBB* distinct from *HBD* (blue dashed line in **g, h**). **i**, Indels at the *HBD* locus measured by deep-seq analysis of sgRNA-1 RNP targeting the β^0^ 4142 site. In all graphs, data are plotted as the mean ± s.d. and analyzed using unpaired two-tailed Student’s *t* tests. Data are representative of three biologically independent replicates.

There are five paralog genes on the human β globin gene cluster consisting of ε globin (*HBE1*), γ globin 2 (*HBG2*), γ globin 1 (*HBG1*), δ globin (*HBD*), and β globin (*HBB*) genes. The sequences and function of *HBB* are most similar to those of *HBD* **(Fig. 2c)**, with 137 of a total of 147 amino acid residues identical^*35*^. We supposed that these restorations in homozygous mutant cells underwent GC from the nearby *HBD* during homology repair. To confirm our hypothesis, we knocked out (KO) the homologous wild-type site of *HBD-4142* in homozygous β^0^ 4142 cells (Donor #1). After 72 hours, the RNP of sgRNA-1 was reintroduced into those cells to target the *HBB-4142* 4 bp deletion site with or without ssODN. Deep sequencing results showed a strong reduction of the restorations compared with the non-*HBD*-KO control **(Fig. 2d)**, either with or without ssODN. Surprisingly, some genotypes of edited cells carried an indel sequence at the β^0^ 4142 site identical to that of *HBD-4142* **(Fig. S2a and b)**, indicating that there was conversion from the edited *HBD-4142* site, which occurred earlier. As far as possible clinically relevant effects, we verified this in multiple β^0^ 4142 heterozygous patient CD34^+^ cells (Donors #2-#5, **Table S2**). Similar results were obtained and suggested that the DNA damage of β^0^ 4142 could be corrected efficiently only through Cas9 protein mixed with sgRNA-1 independent of exogenous DNA templates via GC **(Fig. 2e, Fig. S3a and b)**, as well as the expression of β-globin **(Fig. S3c)**.

Nuclease-mediated DSBs have been reported to stimulate DNA damage responses that can enrich oncogenic cells^*36*^, and to cause large DNA deletions or rearrangements that are difficult to detect by standard amplicon sequencing. We performed clonal analysis of the erythroid progeny of homozygous and heterozygous CD34^+^ HSPCs (Donors #1 and #2) edited by sgRNA-1. First, we genotyped the repaired clones of compound heterozygous Donor #2, which carried a 1 bp substitution (IVS1-1, G>T) 161 bp upstream of the *HBB*-4142 site. Those clones still preserved G>T substitution excluded the possibility of large deletion on the genome due to the Cas9 endonuclease **(Fig. 2f)**. Meanwhile, for homozygous Donor #1, the clones were corrected deletions of TCTT but not carried with upstream specific fragments and downstream base pairs of *HBD* distinct from *HBB* **(Fig. 2g and h)**. Moreover, there were no indels at the *HBD* locus when edited by sgRNA-1 only **(Fig. 2i)**, suggesting the high specificity of sgRNA-1 Collectively, we prove that GC mediated correcting of the β^0^ 4142 deletions independent of exogenous DNA templates.

Homologous recombination repair (HRR) is the most accurate and highly fidelity DNA repair system and involves multiple complex signal transduction pathways. The key protein is the breast cancer susceptibility gene BRCA (BRCA1/2)^*37*^. It interacts with a variety of other repair proteins in a complex system of DNA damage repair, including ATM, PALB2, RAD52 and RAD54^37^. To determine the restoration mechanism of GC, we constructed a homologous recombination defect (HRD) by knocking out BRCA1/2, ATM, PALB2 and RAD52 separately in homozygous CD34^+^ HSPCs (Donor #1) and the HUDEP-2 cell line. The HUDEP-2 cell line (HU4142^del^HBD^-^) carried a 4 bp homozygous deletion at the *HBB-4142* site and a 3 bp homozygous deletion near the *HBD-4142* site **(Table S2)**. Compared with the non-HR^KO^ control, deep sequencing results showed a strong reduction of the restorations electroporated with RNP using sgRNA-1 targeting the *HBB-4142* mutant site with or without ssODN. All samples in the HR-KO group showed a correcting rate below the non-HR^KO^ control analyzed in HUDEP-2 cell lines **(Fig. S4a and b)** and with significantly lower than the restorations when KO *BRCA1/2* and *PALB2* in homozygous Donor #1 **(Fig. S4a and b)**, indicating that the HR-associated proteins worked on GC. Moreover, in HUDEP-2 cells with ssODN **(Fig. S4c)**, we found that some of the repaired cells carried a 3 bp deletion at *HBB-4142* sites, similar to *HBD*^*-*^ **(Fig. S4d and e)**, which proved the repair by autologous GC using *HBD*^*-*^ as an endogenous DNA template.

### Transplantation of human HSPCs into mice

We next investigated whether delivery of Cas9 and sgRNA-1 into homozygous and compound heterozygous β^0^ 4142 CD34^+^ cells can convert the deletion (-TCTT) to normal gene sequences via GC in HSCs *in vivo*. Thus, RNP electroporation with Cas9 and sgRNA-1 was dependent without exogenous DNA templates into CD34^+^ HSPCs from homozygous Donor #1 and heterozygous Donor #5. The edited and unedited CD34^+^ HSPCs were transplanted into the bone marrow (BM) of immunodeficient NOD B6. SCID *Il2rγ*^*−/−*^*Kit*^*W41/W41*^ (NBSGW) mice^38^. To distinguish repair through GC independent of exogenous DNA or HDR via an exogenous DNA template, we designed another template, ssODN-P, which carried 1 bp synonymous mutations in PAM. Subsequently, we extracted bone marrow from the mice for analysis and found similar human marrow engraftment approximately 16 weeks after transplantation with edited and unedited CD34^+^ HSPCs **(Fig. 3a)**. To determine how RNP electroporation with Cas9 of sgRNA-1 targeted the β^0^ 4142 locus and supplemented with 4% glycerol affected differentiation potential and lineage survival, we assessed the human haematopoietic lineages in recipient mouse BM after transplantation. Deep sequencing showed a high rate of indels, and most engrafted human cells maintained approximately 80% indels, similar to the 90% indels observed in the edited cells before transplantation, yielding potent human engraftment while maintaining high restoration frequencies in the repopulating BM cells **(Fig. 3b)**. Flow cytometry using an anti-human CD45 antibody showed that human cells made up approximately 80% of BM in all mice **(Fig. 3c)**. Edited cells showed similar capacities for lymphoid, myeloid, and erythroid engraftment **(Fig. 3d and e)**. Additionally, in BM, β-globin in edited human erythroid cells was elevated by over 30% compared with that in edited human erythroid cells **(Fig. 3f)**. The restorations from 4 bp deletions at *HBB-4142* sites to normal gene sequences maintained 10-15% **(Fig. 3g)**. GC without ssODN increased the fraction of heterozygous Donor #5 from 47% to 60 ± 2.3% (mean ± s.d.) in BM cells compared to unedited control cells **(Fig. 3g)**. The restoration of homozygous Donor #1 was approximately 22-27% restoration observed before transplantation and approximately 12-15% restoration efficiency achieved in repopulating HSCs **(Fig. 3g)**. Notably, almost all the repaired CD34^+^ HSPCs via ssODN-P were eliminated after transplantation, while those cells repaired by GC independent of exogenous template were retained and lineage survival, suggesting that GC have a greater advantage in long-term engrafting HSCs **(Fig. 3g)**.

**Fig. 3.**
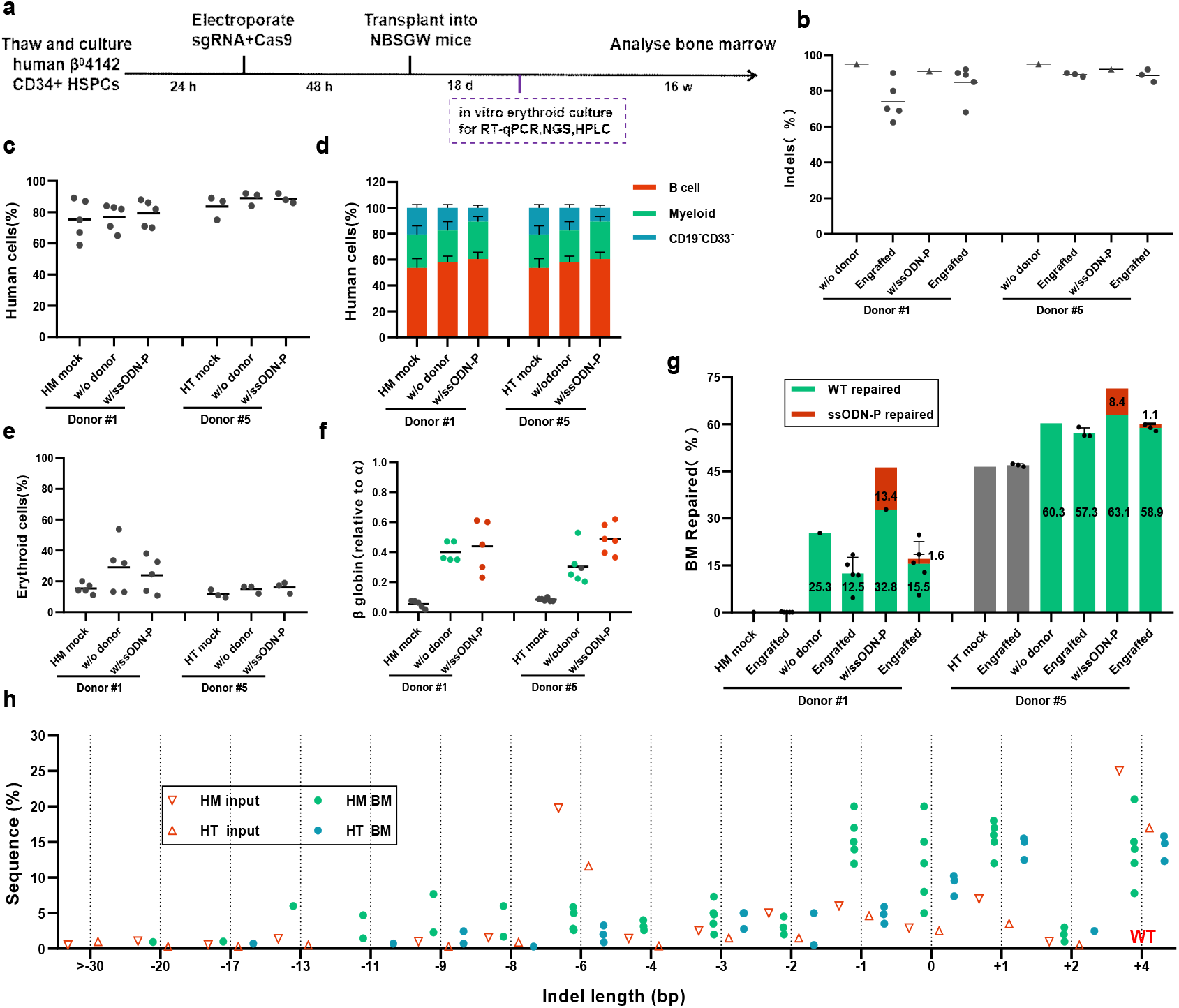
Engraftment of editing β^0^ 4142 CD34^+^ HSPCs after transplantation into immunodeficient mice. CD34^+^ HSPCs from donors with β^0^ 4142 were electroporated with 3NLS Cas9 and sgRNA-1 dependent or independent of exogenous DNA templates. We transplanted 5–8 × 10^5^ treated cells into NBSGW mice via tail-vein injection. Mouse BM was collected and analyzed 16 weeks after transplantation. **a**, Experimental workflow. **b**, In BM, as well as the indel frequencies determined by TIDE analysis. **c**, Engraftment measured by the percentage of human CD45^+^ (hCD45^+^/(hCD45^+^+mCD45^+^)) cells in recipient mouse BM. **d**, Human B cells (hCD19^+^) and myeloid cells (hCD33^+^) as percentages of the hCD45^+^ population in recipient BM. **e**, Human erythroid precursors (hCD235a^+^) as a percentage of human and mouse CD45^−^ cells in recipient BM. **f**, In BM, proportions of β-globin mRNA expression level by RT–qPCR normalized by α-globin. **g**, β^0^ 4142-to-normol editing efficiency in human CD34^+^ cell-derived lineages from recipient BM. **h**. Indel spectrum of input cells from Donor #1 and Donor #5 electroporated with sgRNA-1 before transplantation and BM-engrafted human cells 16 weeks after transplantation. The indel spectrum was determined by deep sequencing analysis. These data comprise 3 mice transplanted from Donor #5 and 5 mice transplanted from Donor #1 with sgRNA-1 edited inputs. Each symbol represents a mouse, and the mean for each group is shown. The median of each group with 3–5 mice in **b, c, e, and f** is shown as a line. Data are plotted as the mean ± s.d. for **d, g** and were analyzed using unpaired two-tailed Student’s *t* tests. Data are representative of three biologically independent replicates.

Moreover, we found that the indel spectrum in repopulating cells was different from that in edited CD34^+^ HSPCs before transplantation **(Fig. 3h)**. In HSPCs edited with Cas9 before transplantation, the second most common indels were 6 bp deletions, comprising 19.9% of alleles in Donor #1 (10.9% in Donor #5) **(Fig. 3h and Fig. S3a-d)**. However, these deletions were significantly absent in the engrafted cells, comprising 3.5% in Donor #1 (1.9% in Donor #5). Consistent with our previous research^18^, large fragment deletions (>-30 bp) were almost absent in the engrafted cells, while the 1 bp insertions and 1 bp deletions were predicted products **(Fig. S3a-d)**. Genetic correction of hemoglobinopathies depends on the fraction of corrected cells that durably repopulate the BM following transplantation. The phenotypic rescue observed following transplantation of varying proportions of edited and unedited mouse BM supports these findings and demonstrates that the editing efficiency achieved in this study substantially exceeds the threshold needed for therapeutic benefit.

### Safety capability evaluation of Cas9 nuclease treatment

The most general concern for genome editing is off-target genotoxicity. To analyze the off-target activity of sgRNA-1 Cas9 RNPs in β^0^ 4142 CD34^+^ cells, 20 potential genomic off-target sites **(Table S3)** with three or fewer mismatches relative to the on-target site and predicted off-target sites within the δ-globin genes (*HBD*)^35^ were identified by a common off-target prediction tool^39^. Then, we performed amplicon deep sequencing^40^ for each of these sites in genomic DNA samples isolated from cells with efficient on-target editing and control cells that were not edited (Cas9 only and mock). Most off-target sites showed no Cas9-dependent indel formation within the limit of detection (∼0.1%), and *HBD* was not detected **(Fig. S5a)**. Thus, at these sites, we did not detect evidence of off-target editing in edited cells compared to unedited cells.

Consistent with intact DNA damage responses electroporated with Cas9: sgRNA-1 RNP in HSPCs, we measured the expression of *CDKN1* (*p21*)-DNA damage response Nuclease-treated cells^*41, 42*^ showed peak levels (4.5-fold higher levels) between treatment 6-10 hours and then decreased after 72-96 hours compared with Mock cells and Cas9 only that had been electroporated without sgRNA **(Fig. S5b)**. Taken together with the repair of GC, it appeared likely that most restorations had been finished in 72 hours when electroporating with Cas9 nuclease at the target site, indicating that this DNA damage response had no impact on HSPCs potential.

### CRISPR/Cas9-mediated GC to correct multiply mutations in *HBB* coding region*s*

*Considering the universality and feasibility of this mechanism, HBD* can be used as a template for GC to repair *HBB* lesions **(Fig. 4a)**. Here, we provide evidence that β-thalassemia of varying genotypes in *HBB* coding region*s, including HBB*-IVS1-1 (G>T), CD17 (A>T), HbE (A>G), CD7172 (+A), and SCD (A>T) mutations^31^, were indeed rescued in GC **(Fig. 4b, Table S2)**. Compared with mock cells, the restoration from mutations to wild-type sequences was approximately 10-15%, with elevated β-globin (relative to α-globin) mRNA levels (mean 37%, compared with 1.8% in unedited mock) **(Fig. 4c and Fig. S6a-d)**.

**Fig. 4.**
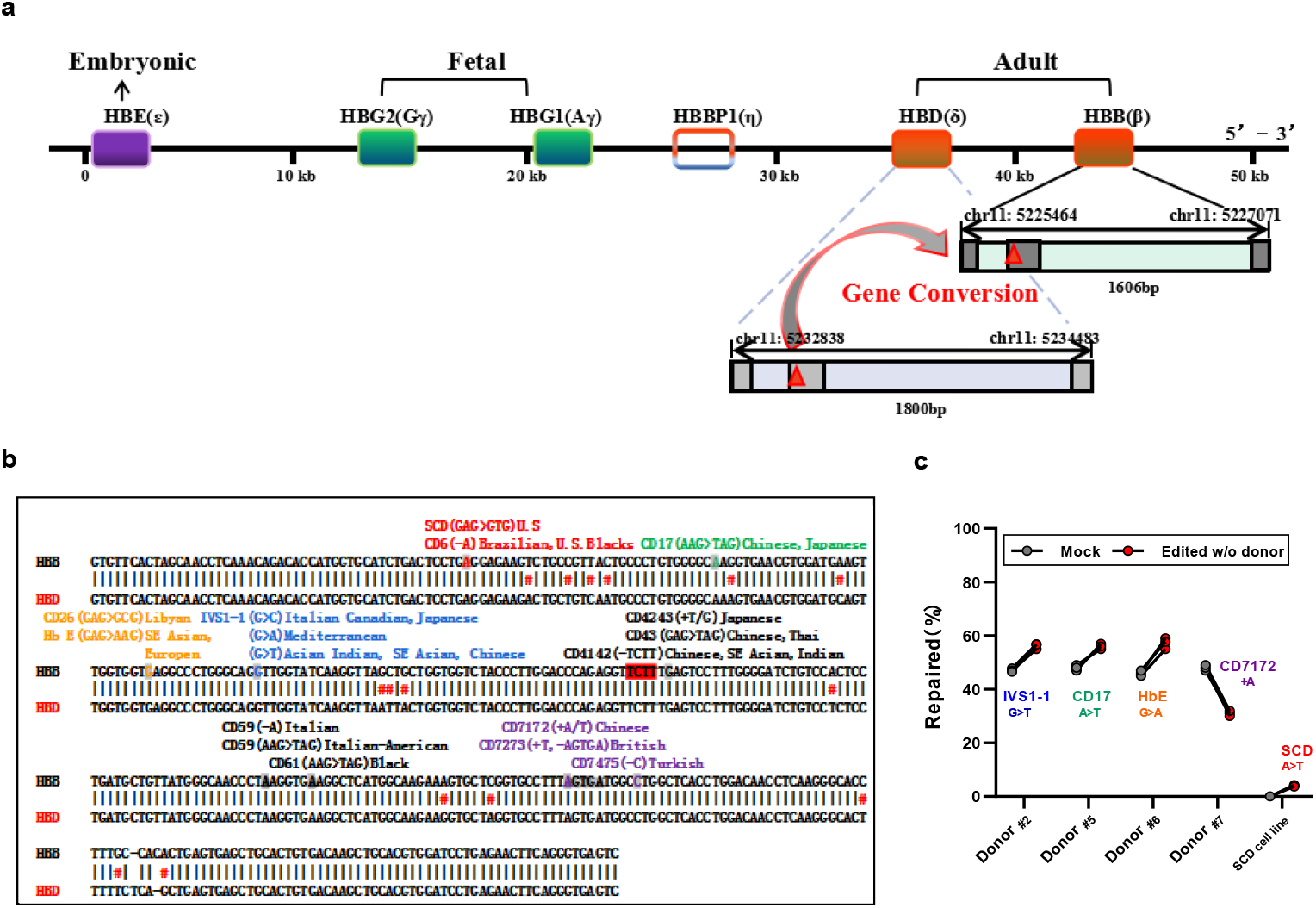
Correcting varying genotypes of β-thalassemias mutations by GC in patient CD34^+^ HSPCs. Efficient correction could be achieved with GC of multiple types mutations in the *HBB* coding regions. **a**. Schematic diagram of the human β-globin gene cluster with expression stages labeled. **b**. The *HBB– HBD* homologous region in the coding region. The vertical gray lines indicate the homologous regions; nonhomologous sequences are indicated by a pound sign (#) and colors it red; positions of multiple types β-thalassemia mutations might be corrected with GC are indicated. They are indicated by different colors. **c**. Efficiency of restoration achieved with GC as measured by deep-seq analysis of RNPs targeting mutations in multiple types β-thalassemia cells (sgRNAs; see details in Table S1). Error bars indicate the standard deviation (*n*=3 replicates).

## Discussion

β-thalassemias is one of the most common autosomal recessive inherited disorders in the world and is caused by more than 300 mutations of the *HBB* gene and characterized by a quantitative deficiency of β-globin chains^31, 32^. With respect to gene therapy for β-thalassemia, significant progress is expected, also considering fundamental insights into globin switching and new technology developments. Programmable endonuclease-mediated genome editing of *BCL11A* enhancer hematopoietic cells leverages NHEJ-mediated genetic disruption and is a novel therapeutic option (NCT 03653247, 03432364, 03745287, 03655678, 02500849 and 03399448). Even though these approaches are promising, they are currently still under deep experimental development and limited to a low number of clinical trials^22, 23^. The γ-subunit differs from its adult counterpart in that HbF has a high ability to bind oxygen more readily from the maternal circulation, physiological differences in oxygen delivery that are important to clinical observation, and patients with hereditary persistence of fetal hemoglobin (HPFH) exhibiting high levels of endogenous HbF might exhibit a clinical status^43^.

In the case of most genetic blood disorders, lasting correction requires HDR-mediated editing of endogenous genes in repopulating HSCs. The therapeutic application of homology-dependent recombination might be especially challenging given the requirement for codelivery of exogenous donor DNA templates with toxicity and insertional mutagenesis, particularly because the edits do not typically confer a selective advantage in the target cell compartment, as is the case for correction of the *HBB* mutations in BM stem cells^4^. Instead of using CRISPR/Cas9 to target disease-causing sites to achieve the goal of directly raising β-globin to balance the excess HBA. To the best of our knowledge, the CRISPR/GC strategy offers the potential for precise endogenous substitution with high product purity to directly convert pathogenic mutations into the wild-type allele^44^ independent of exogenous DNA.

We present for the first time the correction of diseases in human somatic cells by CRISPR/GC. Here, we focused on β^0^ 4142 and some other genetic mutations in *HBB* coding region*s*^45, 46^ *using Cas9 RNP to develop a rapid and extensible gene editing pipeline to introduce mutations into normal human adult HSPCs without supplying DNA repair templates. These reagents efficiently induce non-crossover GC-mediated editing of HBB* mutations in HSPCs with minimal genic off-target activity. Collectively, our study showed that the proportions of each lineage were equivalent in mice that received unedited or edited cells. Edited HSCs from donors with β^0^ 4142 can repopulate the haematopoietic system and generate erythroid cells with a greatly reduced propensity for thalassemia. We describe levels of sequence replacement in long-term engrafting HSCs that may be clinically relevant.

The past few years have witnessed significant progress toward an understanding of the mechanism and its impact on human genome evolution. Whether in an evolutionary or a pathological context, focuses almost exclusively on human diseases caused by GC. Here, we used the most effective RNPs to develop ex vivo editing methods in human HSPCs, achieving up to a high level of replacement. After engraftment in immunocompromised mice for 4 months, sequence conversion at the locus is retained at levels likely to have clinical benefit. It is important to not only develop an approach that is inexpensive and accessible to a wide variety of researchers to edit HSPCs as a potential treatment but also enable investigator-led studies and rapid optimization to address a multitude of genetic diseases, including the Rh blood group antigen genes *RHD* and *RHCE*^47^ and *CCR2* and *CCR5*^48,^ relative to immunologic and cardiovascular diseases.

## Methods

### Ethics Satement

All the studies were conducted following the Department of Hematology, The First Affiliated Hospital of Guangxi Medical University, Guangxi, with review board (IRB) approval. Written informed consent was obtained from all β^4142^-thalassemia patient donors prior to enrollment in the study.

### Culture of CD34^+^ human HSPCs

β-Thalassemia CD34^+^ HSPCs (**Table S2**) were enriched using the Miltenyi CD34 Microbead kit (Miltenyi Biotec). CD34^+^ HSPCs were thawed on day 0 into X-VIVO 15 (Lonza, 04–418Q) seeded and maintained at a density of 0.5–1×10^6^ cells per ml and supplemented with 100 ngml^−1^ human stem cell factor (SCF, R&D), 100 ngml^−1^ human thrombopoietin (TPO, R&D), and 100 ngml^−1^ recombinant human Flt3-ligand (Flt3-L, R&D). HSPCs were electroporated with Cas9 RNP 24 h after thawing and maintained in X-VIVO media with cytokines. For in vitro erythroid maturation experiments, 48 h after electroporation, HSPCs were transferred into erythroid differentiation medium (EDM) consisting of StemSpan serum-free expansion medium II(SFEM II; Stem Cell Technologies) with 1% penicillin/streptomycin (Thermo Fisher Scientific), 3 IU ml^−1^ erythropoietin (Amgen) and 5% human solvent detergent pooled plasma AB. During days 0–11 of culture, EDM was further supplemented with 100 ngml^−1^ human SCF and 5 ngml^−1^ human IL-3 (R&D) as EDM-1. During days 11–18 of culture, EDM had no additional supplements. The enucleation percentage and β-globin induction were assessed on 18 d of erythroid culture.

### Cas9 nuclease purification

We transformed pET-21a^_^3×NLS–Streptococcus pyogenes Cas9 available on Addgene (SpCas9, ID #114365) into BL21 (DE3) chemically competent cells (TransGen Biotech, CD601-02) and grew the cells in LB medium at 37°C 220 rpm until the density reached OD600 = 2-2.5. Cells were induced with 0.2 mM isopropyl β-d-1-thiogalactopyranoside (IPTG) per liter for 16 h at 18°C. Cell pellets were lysed in 20 mM Tris, 10 mM imidazole, 500 mM NaCl, 1.5 M urea pH 8.0 by ultrasonic crushing and centrifuged at 4,000 rpm for 1 h at 4°C. The protein was purified from the cell lysate using nickel-NTA resin and washed with 3 volumes of nickel-NTA buffer. Next, the protein was purified by cation exchange chromatography (SP and heparin column). Endotoxin was controlled by anion exchange chromatography (Q-HP column) (**Table S4**). Subsequently, purified proteins were concentrated and filtered using Amicon ultrafiltration units with a 30-kDa MWCO (Millipore Sigma, UFC903008) and an Ultrafree-MC centrifugal filter (Millipore Sigma, UFC30GV0S) assessed by SDS–PAGE (Sure PAGE, Bis-Tris M00662) to be > 95% pure, and the protein concentration was quantified with the Pierce BCA Protein Assay kit (Thermo Scientific REF23227).

### RNP electroporation

Electroporation was performed using a Lonza 4D Nucleofector (V4XP-3032 for 20 μl Nucleocuvette Strips or V4XP-3024 for 100 μl Nucleocuvettes) according to the manufacturer’s instructions. The modified synthetic sgRNA (2′-*O*-methyl and phosphorothioate modifications at the first three 5′ and 3′ terminal RNA residues) was purchased from Nanjing GenScript Biotech Co., Ltd. HSPCs were thawed 24 hours before electroporation. For 20 μl Nucleocuvette Strips, the RNP complex was prepared by mixing Cas9 (100 pmol) and sgRNA (450 pmol, full-length product reporting method) and incubating for 10-20 min at room temperature immediately before electroporation. HSPCs (1×10^5^ cells) resuspended in 20 μl of P3 solution RNP added to 4% glycerol solution were mixed with and transferred to a cuvette for electroporation with the program EO-100. For 100 μl cuvette electroporation, the RNP complex was made by mixing 500 pmol Cas9 and 3,000 pmol sgRNA. HSPCs (5 M) were resuspended in 100 μl of P3 solution for RNP electroporation as described above. The electroporated cells were resuspended in X-VIVO media with cytokines and changed to SFEMII medium 24 h later for in vitro differentiation. For mouse transplantation experiments, cells were maintained in X-VIVO 15 with SCF, TPO, and Flt3-L for days 0–2 as indicated before infusion.

### Clonal culture of CD34^+^ HSPCs and HUDEP-2 cell lines

Edited CD34^+^ HSPCs were sorted into 150 μl EDM-1 in 96-well round-bottom plates (Nunc) at one cell per well using a Blood Cell Counter. The medium was changed to 500 μl EDM media 7 days later in 24-well flat-bottom plates for further differentiation (Nunc). After 4-7 days of culturing, the cells were collected with sufficient material for DNA isolation with a TIANamp Genomic DNA Kit (DP304, TIANGEN) for genotyping analysis in technical triplicate for genotyping analysis. For HUDEP-2 culture, the medium consisted of SFEM (Stem Cell Technologies) with 1% penicillin/streptomycin, 3 IU ml^−1^ erythropoietin, 50 ngml^−1^ human SCF, 1 μgml^−1^ doxcycline (TAKARA Bio) and 10^−6^ M dexamethasone (Sigma Aldrich).

### Human CD34^+^ HSPC transplantation and flow cytometry analysis

All animal experiments were approved by the East China Normal University Animal Care and Use Committee. CD34^+^ HSPCs were obtained from anonymized β-hemoglobinopathy patients under protocols approved by the IRB of The First Affiliated Hospital of Guangxi Medical University, with the informed consent of all participants, and complied with relevant ethical regulations. *NOD. Cg-Kit*^*W41J*^*Tyr+Prkdc*^*scid*^*Il2rg*^*tm1Wjl*^ (NBSGW) mice were purchased from GemPharmatech Co, Ltd. Non-irradiated NBSGW female mice (4–5 weeks of age) were infused by retro-orbital injection with 0.5–1 M CD34^+^ HSPCs (resuspended in 200 μl DPBS) derived from β-hemoglobinopathy patients. Equal numbers of pre-electroporation CD34^+^ HSPCs were used for experiments comparing in vitro culture for 0, 1, or 2 days following electroporation. Bone marrow was isolated for human xenograft analysis 16 weeks post-engraftment. Serial transplants were conducted using retro-orbital injection of bone marrow cells from the primary recipients. For flow cytometry analysis, bone marrow cells were first incubated with Human TruStain FcX (422302, BioLegend) and TruStain fcX (anti-mouse CD16/32, 101320, BioLegend) blocking antibodies for 10 min, followed by incubation with V450 Mouse Anti-Human CD45 Clone HI30 (560367, BD Biosciences), PE-eFluor 610 mCD45 Monoclonal Antibody (30-F11) (61–0451–82, Thermo Fisher), FITC anti-human CD235a Antibody (349104, BioLegend), PE anti-human CD33 Antibody (366608, BioLegend), APC anti-human CD19 Antibody (302212, BioLegend), and Fixable Viability Dye eFluor 780 for live/dead staining (65-0865-14, Thermo Fisher). Percentage human engraftment was calculated as hCD45^+^ cells/ (hCD45^+^ cells + mCD45^+^ cells) ×100. B-cells (CD19^+^) and myeloid (CD33^+^) lineages were gated on the hCD45^+^ population. Human erythroid cells (CD235a^+^) were gated on the mCD45^−^hCD45^−^ population. For staining with immunophenotype markers of HSCs, CD34^+^ HSPCs were incubated with Pacific Blue anti-human CD34 Antibody (343512, Biolegend), PE/Cy5 anti-human CD38 (303508, Biolegend), APC anti-human CD90 (328114, Biolegend), APC-H7 Mouse Anti-Human CD45RA (560674, BD Bioscience), and Brilliant Violet 510 anti-human Lineage Cocktail (348807, Biolegend). Cells were resuspended in prewarmed HSPC medium. First, we added Hoechst 33342 to a final concentration of 10 μgml^−1^ and incubated the cells at 37°C for 15 min. Then, we added Pyronin Y directly to the cells at a final concentration of 3 μgml^−1^ and incubated them at 37°C for 15 min. After washing with PBS, we performed flow cytometric analysis or cell sorting. Cell sorting was performed on a FACSAria II machine (BD Biosciences).

### Imaging flow cytometry analysis

In vitro differentiated 18 d erythroid cells stained with Hoechst 33342 were resuspended with 150 μl DPBS for analysis with Imagestream X Mark II (Merck Millipore). Well-focused Hoechst-negative single cells were gated for circularity analysis with IDEAS software. Cells with circularity scores above 15 were further gated to exclude cell debris and aggregates. No fewer than 2,000 gated cells were analyzed to obtain a median circularity score.

### Measurement of indel and repaired frequencies

Indel frequencies were measured with cells cultured in SFEM II 5 days after electroporation. Briefly, genomic DNA was extracted using the TIANamp Genomic DNA Kit (DP304, TIANGEN). The *HBB* codon 41/42 functional core was amplified (618 bp) with 2×Hieff Canace^®^ Gold PCR Master Mix (YEASEN), and the corresponding primers were: 4142 F: 5′-GCTTCTGACACAACTGTGTTC-3′; 4142 R: 5′-CCACACTGATGCAATCATTCG-3′ using the following cycling conditions: 98°C for 3 min; 34 cycles of 98°C for 10 s, 6 0°C for 15 s, and 72°C for 15 s; 72°C for 5 min. For deep sequencing, the *HBB* codon 41/42 functional core was first amplified using the following cycling conditions: 98°C for 3 min; 34 cycles of 98°C for 10 s, 63°C for 20 s, and 72°C for 7 s; 72°C for 5 min, with the corresponding primers: 4142NGS-F: 5′-<u>GGAGTGAGTACGGTGTGC</u>AGGAGACCAATAGAAACTGGGC-3′; 4142NGS-R: 5′-<u>GAGTTGGATGCTGGATGGC</u>ACTAAAGGCACCGAGCACT-3′.

Underlined sequences represent adapters for Illumina sequencing. The resulting PCR products were subjected to Sanger sequencing or Illumina deep sequencing, and sequencing traces were imported into TIDE software for indel frequency measurement with a 40 bp decomposition window. After another round of PCR with primers containing sample-specific barcodes and adaptors, amplicons were sequenced for 250 paired-end reads with the MiSeq Sequencing System (Illumina). Frequencies of editing outcomes were quantified using CRISPResso2 software^40^ and collapsed on the basis of mutations in the quantification window. Indels overlapping the spacer sequence were counted as indels, and the number of reads +TCTT was counted as the mock amplicon length for total edit quantification.

### Amplicon deep sequencing

For indel frequencies or off-target analysis with deep sequencing, *HBB* codon 4142 loci or potential off-target loci (*HBD*, in which the amplicon includes homologous genomic sequences) were amplified with corresponding primers first (**Table S3**). After another round of PCR with primers containing sample-specific barcodes and adapters, amplicons were sequenced for 250 paired-end reads with a MiSeq Sequencing System (Illumina). The deep sequencing data were analyzed by CRISPResso software^40^.

### Determination of *HBB* mRNA

Cells were directly lysed into RLT plus buffer (TIANGEN, DP430) for total RNA extraction according to the manufacturer’s instructions provided in the RNeasy Plus Mini Kit. β-globin mRNA expression was internal controled amplifying *HBA* with the following primers: HBA-S_qPCR: GCCCTGGAGAGGATGTTC; HBA-AS_qPCR: TTCTTGCCGTGGCCCTTA; HBB-S: TGAGGAGAAGTCTGCCGTTAC; HBB-AS: ACCACCAGCAGCCTGCCCA. All gene expression data represent the mean of at least three technical replicates. For in vitro differentiation, unless otherwise indicated, *HBB* mRNA levels were measured on 18 d.

### Statistics and reproducibility

We utilized unpaired two-tailed Student’s *t* test, Pearson correlation, and Spearman correlation using GraphPad Prism for analyses, as indicated in the figure legends.

## Acknowledgment

We are indebted to the Xiangya Hospital clinical staff, especially β-thalassemia patients who contributed samples for this study. We thank the support of the ECNU Public Platform for innovation (011). We thank BRL Medicine Inc., W. Li for assistance with patient recruitment.

## Author contribution

Y.W., D.L., F.Y. designed the experiments and analyzed the data; D.L. and F.Y. wrote the manuscript. F.Y. performed all experiments in human CD34^+^ HSPC, RNP editing, CD34^+^ HSPC transplant, and engraftment analysis. G.L. and Y.Z. assisted with flow cytometry. Assisted by S.Q. and J.P. produced Cas9 proteins; Y.Y. and Y.L. provided the HPSCs and analyzed the data. D.L. and Y.W. conceived and supervised the study.

