## Supplemental Figure S1-S6, Table S1-S4 for "Efficient repair of human homozygous genetic mutation by CRISPR/Cas9 mediated interlocus gene conversion"

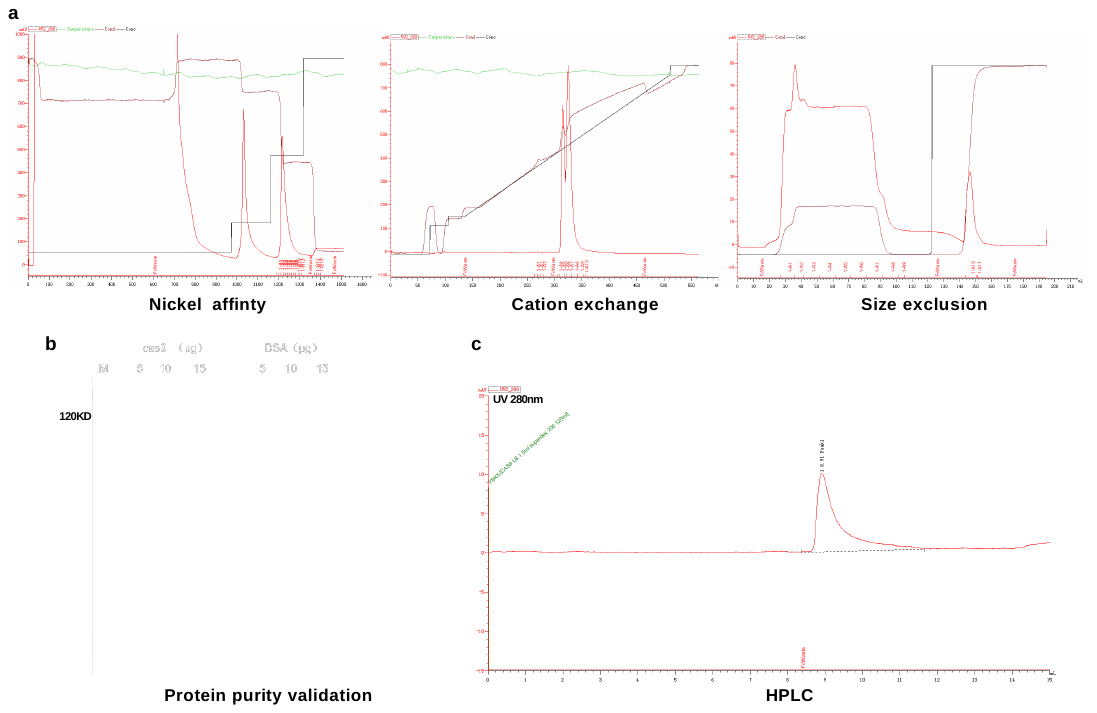
 Fig. **S1| Purification of the 3NLS-Cas9 protein. a**, Purification strategy and profiles. The 3NLS-Cas9 protein was purified using nickel affinity, mono S cation exchange, and size exclusion columns. Desired fractions from 100% imidazole elution (nickel column tube A1-B11), salt gradient elution (tube A4-A9), and size exclusion (tube A1–A7) were collected. **b**, Protein purity and concentration validation. Protein purity was determined using SDS PAGE gel staining and BAS. Total clear protein lysate and ion exchange (IEX) samples and fractions were loaded on SDS–PAGE gels and stained with GelCode Blue to check the purity. **c**, After immobilized metal affinity chromatography and IEX, the protein purity was estimated to be more than 95%. Protein purification was performed once.

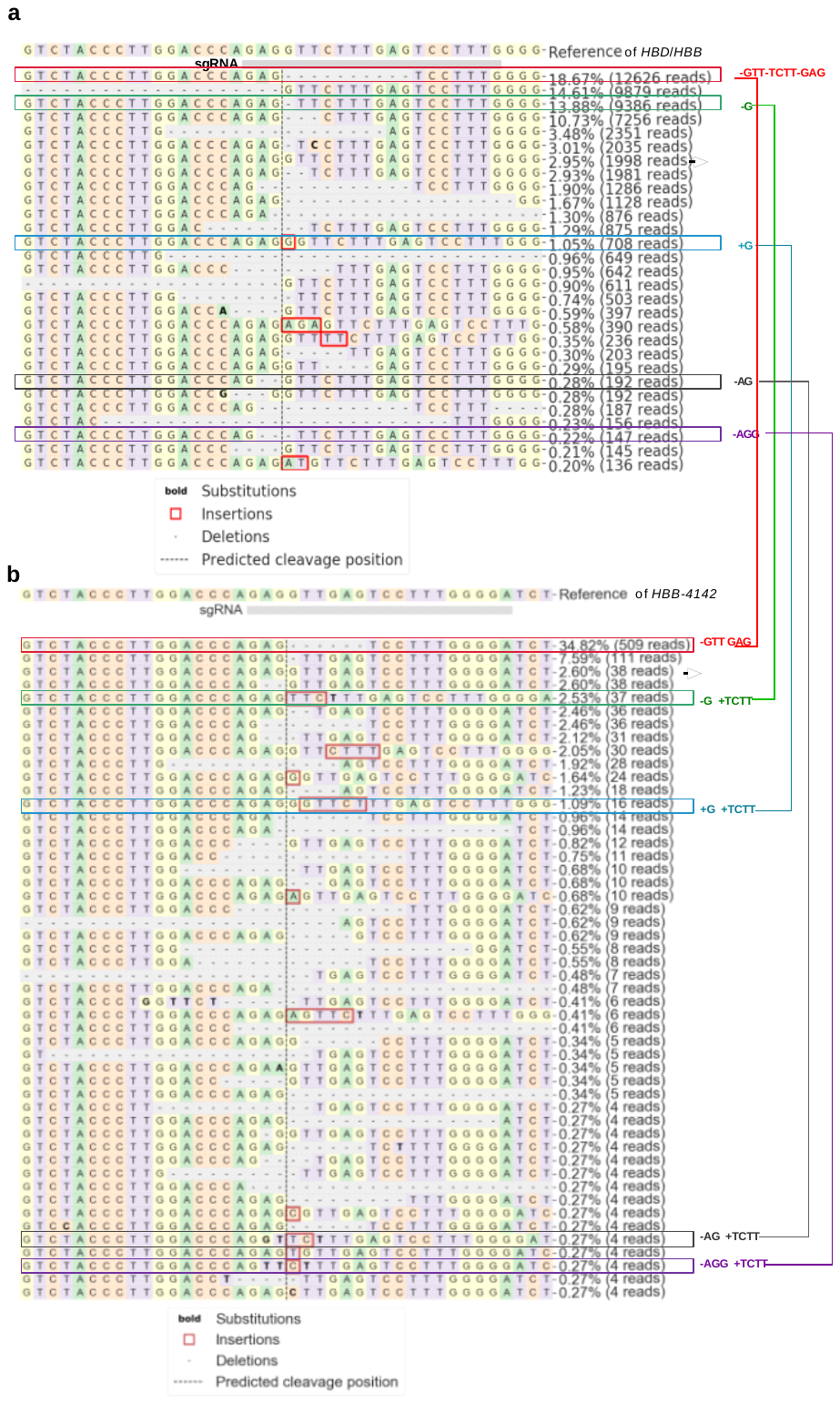

Fig. S2**|GC in homozygous mutant cells via the nearby HBD during homology repair.** **a**, Summary of the most frequent indels by deep sequencing following Cas9 RNP *HBD-4142* editing of homozygous β^0^ 4142 CD34^+^ HSPCs (*HBD*^KO^). **b**, Summary of deep sequencing data derived from Cas9 RNP (coupled with sgRNA-1)-edited homozygous β^0^ 4142 CD34^+^ HSPCs (*HBD*^KO^). The arrows indicate unedited alleles. The same genotypes with measurable conversions were in the same color.

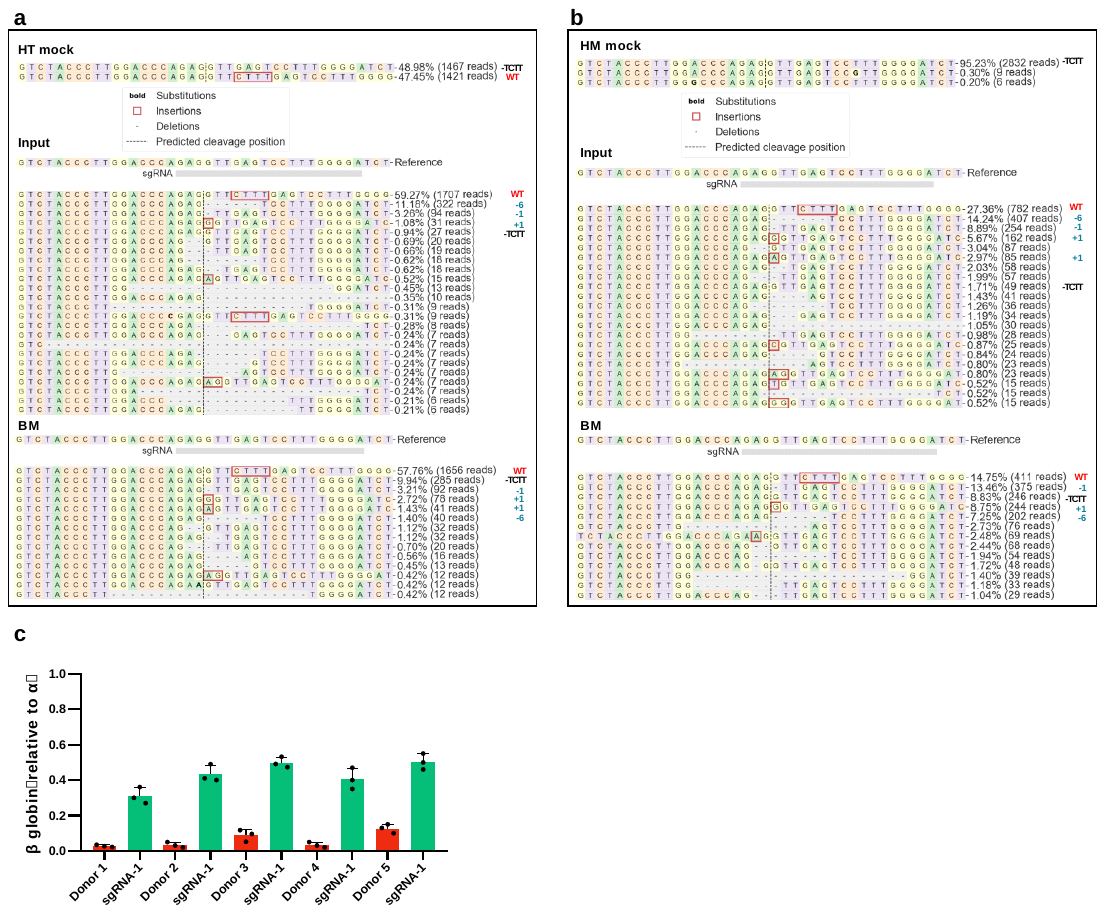
Fig. S3**|Indel spectra of engrafted BM correcting β^0^ 4142 through GC. a**, **b**, Summary of the most frequent indels by deep sequencing of input cells and corresponding bone marrow cells from the primary recipient. (**a**) Restoration by deep sequencing following Cas9 RNP editing of Donor #5 (HT). (**b**) Restoration by deep sequencing following Cas9 RNP editing of Donor #1 (HM). HT, HM mock, heterozygote and homozygous β^0^ 4142-thalassemia patient CD34^+^ HSPCs; Input, deep sequencing following Cas9 RNP editing of patient CD34^+^ HSPCs before transplantation; BM, engrafted edited CD34^+^ HSPCs analyzed 16 weeks after transplantation. -TCTT indicates unedited allele; WT remarks in red indicate normal wild-type alleles; Green indicates other most frequent indels. **c**, RT–qPCR analysis showed that the expression of β-globin was rescued in 4142 deletion HSPCs edited with sgRNA-1 that underwent GC. Error bars indicate the standard deviation (*n*= 3 replicates).

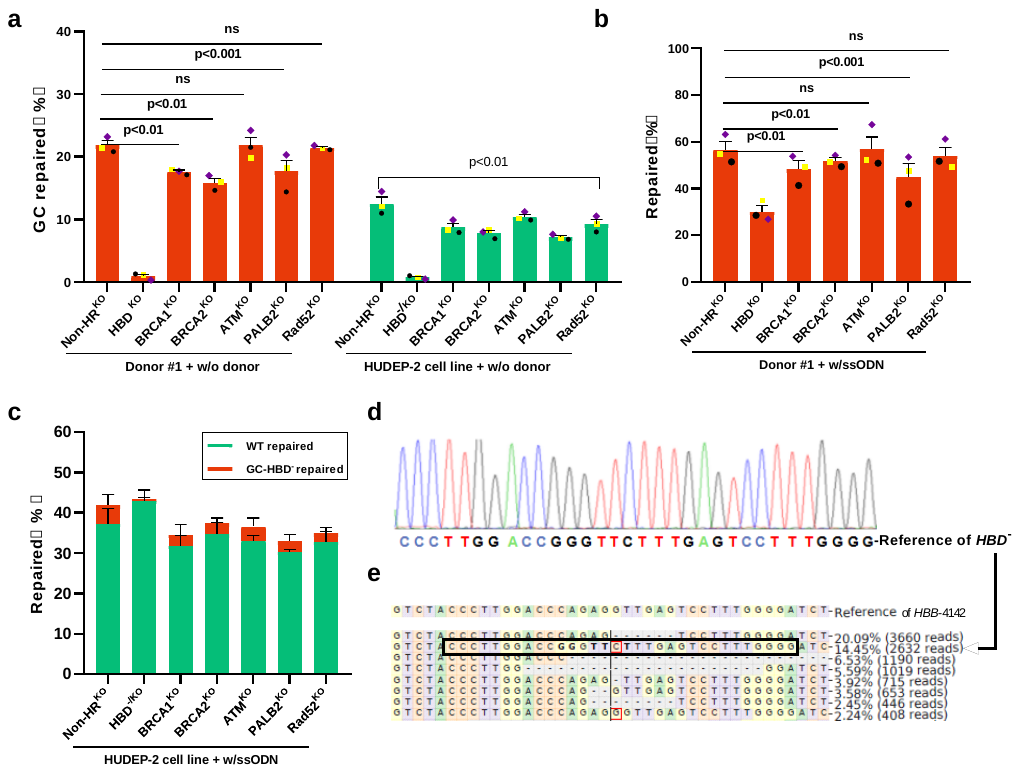

Fig. S4**|GC- and HR-associated proteins** **a-c**, Comparing homologous recombination (HR) and GC when inhibiting HR-related genes (sgRNAs; see details in Table S1) in the homozygous Donor #1 and HUDEP-2 cell lines. After 72 hours, the cells were electroporated with RNP using sgRNA-1 targeting the mutant site with or without exogenous DNA templates. Data are plotted as the mean ± s.d. and analyzed using unpaired two-tailed Student’s *t* tests. NS, not significant. Data are representative of three biologically independent replicates. **d**, The *HBD* genotype of the HUDEP-2 cell line. **e**, Summary of the most frequent indels by deep sequencing of sgRNA-1 RNP targeting the *HBB-4142* site in the HUDEP-2 cell line. Genotype of *HBD^-^* remarks in black box indicates GC restorations;

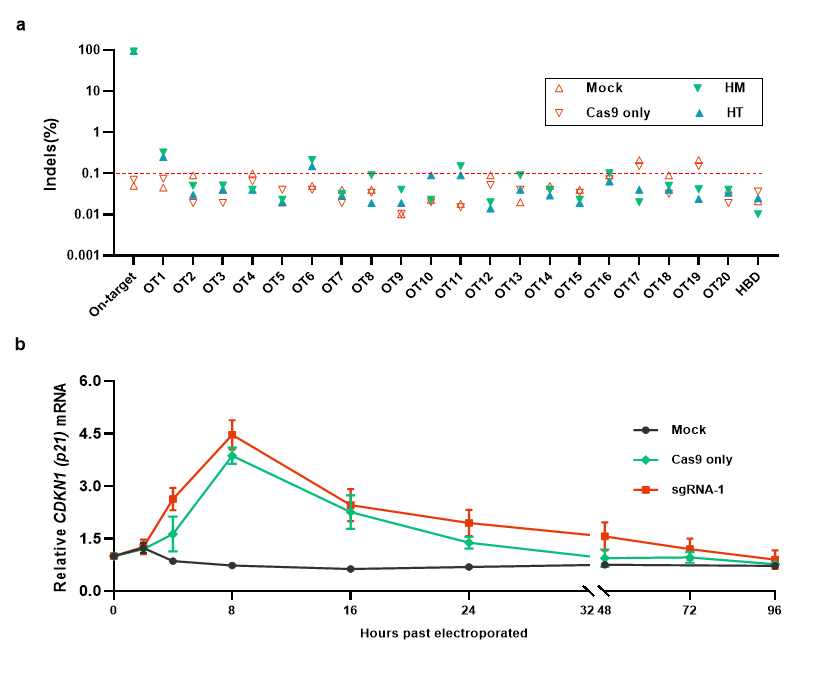

Fig. S5**|Off-target analysis edited by Cas9 RNP targeting sgRNA-1 conversion of *HBB*-4142 deletion** **to normal in homozygous and heterozygous patient donors. a.** RT–qPCR analysis of *p21* expression after gene editing. Relative expression to β-actin is shown. Error bars indicate the standard deviation (*n*= 3 replicates). **b**. Using the CasOFFinder tool, 21 potential genomic off-target sites with 3 or fewer mismatches to the on-target sgRNA-1 sequence were evaluated by amplicon deep sequencing. The on-target sequence is at the *HBB-4142* site. The dotted line at 0.1% denotes the sensitivity of deep sequencing to detect indels.

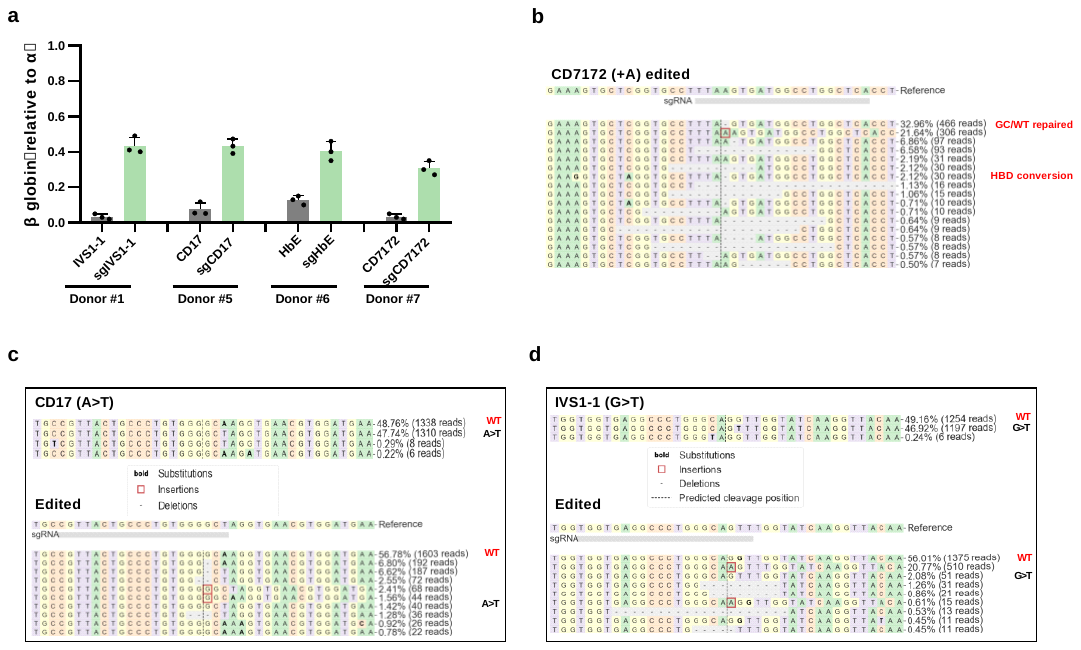

Fig. S6**|GC to correct multiple [types](https://cn.bing.com/dict/search?q=types&FORM=BDVSP6&cc=cn) of [mutation](https://cn.bing.com/dict/search?q=mutation&FORM=BDVSP6&cc=cn)s in the [coding](https://cn.bing.com/dict/search?q=coding&FORM=BDVSP6&cc=cn) [region](https://cn.bing.com/dict/search?q=region&FORM=BDVSP6&cc=cn) *HBB.* a.** β-globin expression by RT–qPCR analysis in erythroid cells in vitro differentiated from RNP-edited β-thalassemia patient donors. Error bars indicate the standard deviation (*n*= 3 replicates). **b.** Summary of the restorations by deep sequencing following Cas9 RNP editing of CD7172. The genotypes with measurable conversion from *HBD* are labeled in red. **c.** Summary of the restorations by deep sequencing following Cas9 RNP editing of CD17. **d.** Summary of the restorations by deep sequencing following Cas9 RNP editing of IVS1-1. WT marked in red indicates normal wild-type allele; mutation sites marked in black indicate unedited allele.

| **Table S1. Sequence of sgRNAs and DNA templates** | | |
| --- | --- | --- |
| **sgRNAs** | **5'-3'** | **Target regions** |
| sgRNA-1 | UCCCCAAAGGACUCAACCUC | Targeting HBB-4142 |
| sgRNA-2 | CCCCAAAGGACUCAACCUCU | Targeting HBB-4142 |
| sgRNA-3 | GACCCAGAGGUUGAGUCCUU | Targeting HBB-4142 |
| sgRNA-4 | ACCCAGAGGUUGAGUCCUUU | Targeting HBB-4142 |
| sgRNA-5 | CCCAGAGGUUGAGUCCUUUG | Targeting HBB-4142 |
| sgRNA-6 | GGACUCAACCUCUGGGUCCA | Targeting HBB-4142 |
| sgRNA-7 | GACUCAACCUCUGGGUCCAA | Targeting HBB-4142 |
| HBD4142 ^KO^ | CAAAGGACUCAAAGAACCUC | Targeting HBB/HBD WT4142 sites |
| HU-HBD^-^ | CCCAAAGGACUCAAAGAACC | Targeting HBD^-^ mutant site in HUDEP-2 cell line |
| BRCA1^KO^ | AGCAGUAUUUCAUUGGUACC | Targeting BRCA1 gene |
| BRCA2^KO^ | UCUACUAUAUUAGAAGAAUC | Targeting BRCA2 gene |
| ATM^KO^ | CCUUUAGGGCAGCUGAUAUU | Targeting ATM gene |
| PALB2^KO^ | CUUGGAUGAUGAUGCUUUCA | Targeting PALB2 gene |
| Rad54^KO^ | UAAGGCUUUUAAGUCUGUGU | Targeting Rad54 gene |
| sgIVS1-1 | UGGUGAGGCCCUGGGCAGUU | Targeting IVS I-1 G>T in Donor #2 |
| sgCD17 | CGUUACUGCCCUGUGGGGCU | Targeting CD17 A>T in Donor #5 |
| sgHbE | CGUGGAUGAAGUUGGUGGUA | Targeting HbE G>A in Donor #6 |
| sgCD7172 | UGAGCCAGGCCAUCACUUAA | Targeting CD7172 +A in Donor #7 |
| sgSCD | GUAACGGCAGACUUCUCCAC | Targeting Sickle mutation A>T in SCD cell line |
| **DNA templates** | **5'-3'** | |
| ssODN | TCCCACCCTTAGGCTGCTGGTGGTCTACCCTTGGACCCAGAGGT**TCTT**TGAGTCCTTTGGGGATCTGTCCACTCCTGATGCTGTTATGGG | |
| ssODN-R | CCCATAACAGCATCAGGAGTGGACAGATCCCCAAAGGACTCA**AAGA**ACCTCTGGGTCCAAGGGTAGACCACCAGCAGCCTAAGGGTGGGA | |
| ssODN-P | TCCCACCCTTAGGCTGCTGGTGGTCTACCCTTGGAC**A**CAGAGGT**TCTT**TGAGTCCTTTGGGGATCTGTCCACTCCTGATGCTGTTATGGG | |
| ss-AmC6 | TCCCACCCTTAGGCTGCTGGTGGTCTACCCTTGGACCCAGAGGT**TCTT**TGAGTCCTTTGGGGATCTGTCCACTCCTGATGCTGTTATGGG | |
| ss-AmC12 | TCCCACCCTTAGGCTGCTGGTGGTCTACCCTTGGACCCAGAGGT**TCTT**TGAGTCCTTTGGGGATCTGTCCACTCCTGATGCTGTTATGGG | |
| Note： | | |
| Letters in Green represent 4 bp deletion sites | | |
| Lerrer in Red represents synonymous mutations in PAM of ssODN-P. | | |
| ss-AmC6 represents 5′ modifications including an amine group with a C6 linker (AmC6) | | |
| ss-AmC12 represents 5′ modifications including an amine group with a C12 linker (AmC12) | | |

| **Table S2. Genotypes of β-thalassemia patient donors and cell lines** | | | |
| --- | --- | --- | --- |
| **Donor ID** | **Genotype information** | | **Sequence information** |
| 1 | β^4142^β^4142^ | β^0^ | Homozygous for Codon 41/42 (-TCTT)  CD4142: TGGTGGTCTACCCTTGGACCCAGAGGT- - - -TGAGTCCTTTGGGGATCTGTCCACTCCTGAT |
|  |  | β^0^ |  |
| 2 | β^4142^β^IVS I-1^ | β^0^ | Compound heterozygous for Codon 41/42 (-TCTT) and IVS I-1 G>T  CD4142: TGGTGGTCTACCCTTGGACCCAGAGGT- - - -TGAGTCCTTTGGGGATCTGTCCACTCCTGAT IVS I-1 G>T: GGATGAAGTTGGTGGTGAGGCCCTGGGCAG**T**TTGGTATCAAGGTTACAAGACAGGTTT |
|  |  | β^0^ |  |
| 3 | β^4142^β^-28^ | β^0^ | Compound heterozygous for Codon 41/42 (-TCTT) and -28 (A>G) (promoter TATA box) CD4142: TGGTGGTCTACCCTTGGACCCAGAGGT- - - -TGAGTCCTTTGGGGATCTGTCCACTCCTGAT  -28: GGAGGGCAGGAGCCAGGGCTGGGCATA**G**AAGTCAGGGCAGAGCCATCTATTGCTTACATTTG |
|  |  | β^+^ |  |
| 4 | β^4142^β^-29^ | β^0^ | Compound heterozygous for Codon 41/42 (-TCTT) and -29 (A>G) (promoter TATA box) CD4142: TGGTGGTCTACCCTTGGACCCAGAGGT- - - -TGAGTCCTTTGGGGATCTGTCCACTCCTGAT -29: GGAGGGCAGGAGCCAGGGCTGGGCAT**G**AAAGTCAGGGCAGAGCCATCTATTGCTTACATTTG |
|  |  | β^+^ |  |
| 5 | β^4142^β^CD17^ | β^0^ | Compound heterozygous for Codon 41/42 (-TCTT) and Codon 17 (A>T; AAG>TAG; Lys>stop codon)  CD4142: TGGTGGTCTACCCTTGGACCCAGAGGT- - - -TGAGTCCTTTGGGGATCTGTCCACTCCTGAT CD17: GAAGTCTGCCGTTACTGCCCTGTGGGGC**T**AGGTGAACGTGGATGAAGTTGGTGGTGAGGCCC |
|  |  | β^0^ |  |
| 6 | β^Hb E^β^-28^ | β^+^ | Compound heterozygous for Hb E (G>A; GAG>AAG) and -28 (A>G) (promoter TATA box) Hb E: GGCAAGGTGAACGTGGATGAAGTTGGTGGT**A**AGGCCCTGGGCAGGTTGGTATCAAGGTTACA -28: GGAGGGCAGGAGCCAGGGCTGGGCATAGAAGTCAGGGCAGAGCCATCTATTGCTTACATTTG |
|  |  | β^+^ |  |
| 7 | β^7172^β^4142^ | β^0^ | Compound heterozygous for Codon 7172 (+A; frameshift mutation) and Codon 41/42 (-TCTT)  CD4142: TGGTGGTCTACCCTTGGACCCAGAGGT- - - -TGAGTCCTTTGGGGATCTGTCCACTCCTGAT CD7172: CTCATGGCAAGAAAGTGCTCGGTGCCTTT**(A)**AGTGATGGCCTGGCTCACCTGGACAACCT |
|  |  | β^0^ |  |
| n/a | HU4142^del^HBD^-^ | β^0^ | Homozygous HUDEP-2 cell line for Codon 41/42 (-TCTT), Homozygous for HBD indels (-CAGA +G) CD4142: TGGTGGTCTACCCTTGGACCCAGAGGT- - - -TGAGTCCTTTGGGGATCTGTCCACTCCTGAT  HBD^-^: CTGGTGGTCTACCCTTGGACC**- - G -** GGTTCTTTGAGTCCTTTGGGGATCTGTCC |
|  |  | β^0^ |  |
| n/a | SCD Cell line | β^s^ | Homozygous SCD cell line for SCD (A>T; GAG>GTG) Sickle mutation  SCD: AAACAGACACCATGGTGCATCTGACTCCTG**T**GGAGAAGTCTGCCGTTACTGCCCTGTGGGGC |
|  |  | β^s^ |  |

| **Identified by Cas-OFFinder** | | **GRCh38 reference genome** | | | | **Primers for off-target detection** |
| --- | --- | --- | --- | --- | --- | --- |
| **Target** | **Gene Alias** | **HOMER Annotation** | **chromosome** | **Mismatches** | **Aligned sequence at site** |  |
| On target | HBB |  | chr11 | N/A | TCCCCAAAGGACTCAACCTC | N/A |
| OT1 | ZNF385D-AS2-UBE2E2-AS1 | intergenic | chr3 | 3 | ATTCCAAAGGACTCAACCTC | F- **GGAGTGAGTACGGTGTGC**ACCTTATAGTCGCTGCTTGT R- **GAGTTGGATGCTGGATGG**TACTCTGCTATTTGAACATC |
| OT2 | RP5-1119A7.17 | exon | chr22 | 3 | TCCCCAGAGAGCTCAACCTC | F- **GGAGTGAGTACGGTGTGC**TGCTGACCTGGGAAAATAA R- **GAGTTGGATGCTGGATGG**AGGTGACTGAAGGTCAAGGT |
| OT3 | BMPER | intron | chr7 | 4 | TCCAGAAAGGACCTAACCTC | F- **GGAGTGAGTACGGTGTGC**CTCCCCCTTATTTCCAAGTG R- **GAGTTGGATGCTGGATGG**CTGCTAACAGTGGCAGAGCT |
| OT4 | MOCS1-RP11-552E20.1 | intergenic | chr6 | 4 | TCTAGAGAGGACTCAACCTC | F- **GGAGTGAGTACGGTGTGC**GCTCTGTCCACTGCTAAGTT R- **GAGTTGGATGCTGGATGG**ATCTAGAGAGGCTCCAAGGA |
| OT5 | SMYD4 | exon | chr17 | 4 | CCACCAAAAGAGTCAACCTC | F- **GGAGTGAGTACGGTGTGC**AGGCCATAATTAAGTGTCTT R- **GAGTTGGATGCTGGATGG**TACCACAGGACAGAATGTCC |
| OT6 | RNU6-649P-AC107057.1 | intergenic | chr2 | 4 | ACCCCAAAAGACACAATCTC | F- **GGAGTGAGTACGGTGTGC**CTTTCTAAAGATAAAAATGAG R- **GAGTTGGATGCTGGATGG**ATACAGACAATCAAATTCAAC |
| OT7 | SNORA25-RP11-16D22.2 | intergenic | chr13 | 4 | TTTCCAAAGGACACAAACTC | F- **GGAGTGAGTACGGTGTGC**GAGAAGAACTTGGTACGGTC R- **GAGTTGGATGCTGGATGG**ACTCTGACAGGACTCAGTAT |
| OT8 | TTTY2B | intron | chrY | 2 | TCCCCAAAAGTCTCAACCTC | F- **GGAGTGAGTACGGTGTGC**GCTCTTACAGCTGTCAGTCT R- **GAGTTGGATGCTGGATGG**AGAGTGGATTCCAGTGACAA |
| OT9 | TTTY2 | intron | chrY | 2 | TCCCCAAAAGTCTCAACCTC | F- **GGAGTGAGTACGGTGTGC**GCTCTTACAGCTGTCAGTCT R- **GAGTTGGATGCTGGATGG**AGAGTGGATTCCAGTGACAA |
| OT10 | MAP2 | exon | chr2 | 4 | TCTCCAGTGAACTCAACCTC | F- **GGAGTGAGTACGGTGTGC**CGTGGTCTCAATTCTAGATC R- **GAGTTGGATGCTGGATGG**ACTCATCATGAACTCGGGTC |
| OT11 | FAM117A-FAM117A/KAT7 | intergenic | chr17 | 4 | TTCCCAAACTAGTCAACCTC | F- **GGAGTGAGTACGGTGTGC**CTCAGAGGAACCCGTTCCTA R- **GAGTTGGATGCTGGATGG**AGTCAGTCTTGCCCACTATA |
| OT12 | Gap-AL354822.1 | intergenic | chrUn_GL000218v1 | 4 | GCCCCCAAGGACCCAAACTC | F- **GGAGTGAGTACGGTGTGC**AAGGTCCCTCAGTGGCAGTT R- **GAGTTGGATGCTGGATGG**CTTCCTCCTGTTCTTGGTGT |
| OT13 | GEMIN8-GLRA2 | intergenic | chrX | 4 | TCCCCATAGAAATCAACATC | F- **GGAGTGAGTACGGTGTGC**TGTCAGTGGTTTGAAGTTCC R- **GAGTTGGATGCTGGATGG**TCACAATCCGGCTGAGAGGA |
| OT14 | BARX2-RP11-237N19.3 | intergenic | chr11 | 3 | TCCTCATTGGACTCAACCTC | F- **GGAGTGAGTACGGTGTGC**CCAAGCTGAAATAAAGATGC R- **GAGTTGGATGCTGGATGG**TAGAAGACTAGAACCTGTTG |
| OT15 | AC006227.1-AC096669.1 | intergenic | chr2 | 4 | CCCTCAAAGGTCACAACCTC | F- **GGAGTGAGTACGGTGTGC**GCTCAGAATAAATAGGATACA R- **GAGTTGGATGCTGGATGG**GTCTAGGACCCAGGTCCTTAC |
| OT16 | TNIK | intron | chr3 | 3 | TTCCCAAAGTACTCAGCCTC | F- **GGAGTGAGTACGGTGTGC**TCACTGTCTATATTCTGTCT R- **GAGTTGGATGCTGGATGG**CCAGAGGTTTTGTATGTGCC |
| OT17 | RN7SKP256-AL591034.1 | intergenic | chr6 | 4 | TTCCCAAAGGACAAAACTTC | F- **GGAGTGAGTACGGTGTGC**ATGCAGATGGCTGTTGCATT R- **GAGTTGGATGCTGGATGG**CCTAGTCTTCCAGTCTCATT |
| OT18 | LINC00700-RP11-69C17.2 | intergenic | chr10 | 4 | GGCACAAAGAACTCAACCTC | F- **GGAGTGAGTACGGTGTGC**TCAGTGCTGAGTCATTTGAC R- **GAGTTGGATGCTGGATGG**GTCACATGCATATTTACGCC |
| OT19 | ATP2B4 | intron | chr1 | 3 | TCCCCAAAAGACTTAACCCC | F- **GGAGTGAGTACGGTGTGC**ATGTCTCTCTTTATCCCTGC R- **GAGTTGGATGCTGGATGG**GAAACACATACCCCCAGACC |
| OT20 | C18orf63 | intron | chr18 | 4 | ACCCCAAAGGACCAAAACTC | F- **GGAGTGAGTACGGTGTGC**TTAGGAAGGGTCCCTCGTAG R- **GAGTTGGATGCTGGATGG**CGTTTCTGTCCCAGGATTTC |
| OT21 | HBD | exon | chr11 | 4 | CAAAGGACTCAAAGAACCTC | F- **GGAGTGAGTACGGTGTGC**GTCCCTTGGGCTGTTTTCCT R- **GAGTTGGATGCTGGATGG**CACTCAGCTGAGAAAAAG |

| **Table S4. Buffer for protein purification** | | |
| --- | --- | --- |
| Nickel-NTA column buffer | Buffer A | 20 mM TRIS + 500 mM NaCl + 20 mM imidazole, pH 8.0 |
|  | Buffer B | 20 mM TRIS, 250 mM NaCl, 500 mM Imidazole, 10% glycerol, pH 8.0 |
| SP, Heparin column buffer | Buffer C | 20 mM HEPES +10% glycerol, pH 7.5 |
|  | Buffer D | 20 mM HEPES+1 M NaCl+10% glycerol, pH 7.5 |
|  | Buffer E | 20 mM HEPES + 250 mM NaCl + 1 mm EDTA +10% glycerol, pH 7.5 |
|  | Buffer F | 20 mM HEPES +10% glycerol, pH 7.5 |
|  | Buffer G | 20 mM HEPES +1 M NaCl+10% glycerol, pH 7.5 |
| Q-HP column buffer | Buffer C | 20 mM HEPES +10% glycerol, pH 7.5 |
|  | Buffer D | 20 mM HEPES+1 M NaCl+10% glycerol, pH 7.5 |
